# CellOS: Learning a World Model of Cellular State through Joint Embedding Prediction

**DOI:** 10.64898/2026.06.18.733163

**Authors:** Qi Zhou, Yuan Le, Xiaoning Qi, Hanshuo Lu, Shaole Chang, Yunjia Wu, Haoran Wang, Ruoyao Ran, Xin Li

## Abstract

Foundation models learned from single-cell transcriptomes are central to the prospect of AI virtual cell that can represent, query and predict cellular state. However, most current single-cell foundation models learn from a single view of gene expression and are optimized primarily through reconstruction or next-token prediction. As a result, they capture expression abundance but cannot explicitly reconcile complementary views of cellular state. Here we present CellOS, a multi-view foundation model that learns cellular representations from paired expression and perception views. CellOS integrates complementary views through a scalable three-stage training strategy that combines causal cell-sentence language modelling, function-preserving dense-to-mixture-of-experts expansion and latent-space alignment via an LLM-JEPA objective. Using this framework, we trained a 12-billion-parameter model on 390.5 million single-cell transcriptomes. Across diverse benchmarks spanning cell-state annotation, batch integration and perturbation-response prediction, CellOS consistently outperformed state-of-the-art single-cell foundation models. Together, these results suggest that predictive alignment between complementary cellular views provides a scalable path toward representation-centric cellular world models and transferable AI virtual cells.

## 1 Introduction

Cells execute the programs that underlie development, homeostasis, immunity and disease. Single-cell RNA sequencing (scRNA-seq) made cell state measurable at transcriptome-wide scale, resolving individual cells across tissues, organisms, developmental stages and perturbations [1]. The resulting atlases, including the Human Cell Atlas [2] and CZ CELLxGENE [3], now contain hundreds of millions of profiles. This scale creates both a biological opportunity and a computational challenge: rather than only cataloguing cell types after measurement, we can ask whether models can learn the rules that organize cellular state and use them to predict new states.

This question has driven single-cell biology toward foundation models and, more broadly, toward the artificial intelligence virtual cell (AIVC). As in language and vision, these models are pretrained on broad corpora and then adapted to many tasks, including annotation, integration, gene-network analysis and perturbation modelling [4–7]. The AIVC vision asks for a model that provides a universal representation of cellular state, can be queried through downstream instruments and can ultimately support in silico reasoning about cellular behaviour [8, 9]. Such tasks depend on the representation quality of foundation models: a useful cellular representation must preserve the biological variables that distinguish one state from another, including rare and context-dependent signals, while remaining scalable enough to learn from atlas-scale data.

The dominant language-inspired formulation represents each cell as a cell sentence. Genes are emitted as tokens, commonly ordered or weighted by within-cell expression, and a transformer is trained to reconstruct masked values or predict subsequent tokens [4, 5, 10–12]. This strategy has been remarkably productive. Geneformer showed that rank-value encoding of approximately 30 million transcriptomes can transfer to network-biology tasks [4]; scGPT and scFoundation extended the field toward generative and multi-omic settings at larger scale [5, 6]; Cell2Sentence and C2S-Scale demonstrated that gene-order sentences can make transcriptomes accessible to large language models at increasing scale [10, 13]; and TranscriptFormer showed that generative atlas training across species can recover cell-type, developmental and phylogenetic structure without explicit supervision [7]. These successes show that gene order captures part of cellular state. Yet the same paradigm also exposes what it leaves implicit: a cell sentence usually gives the model one expression-derived ordering of genes, and the objective usually asks the model to recover input-space tokens or values.

Despite this progress, current cell-sentence models remain limited in two ways. First, expression magnitude is not the same as biological informativeness. A highly expressed housekeeping gene can dominate a within-cell ranking while carrying little state-specific signal, whereas a transcription factor or stress-response gene expressed at a moderate level may be highly diagnostic if that value is rare in the population. The relevant quantity is therefore not only the expression value inside one cell, but where that value lies in the global distribution of the same gene. We formalize this population context with a population-relative surprisal score: for gene *g*, each cell-specific expression value is compared with the empirical distribution of that gene across the pretraining corpus, assigning higher information content to values that are unexpectedly elevated relative to the population background. Existing corpus-normalized rankings approximate this intuition, but they collapse magnitude and population rarity into a single order; they do not expose information content as a separate view of the cell. Second, input-space objectives spend capacity on reconstructing tokens or counts, even when the desired object is a robust cell-level representation. In vision and language, joint-embedding predictive architectures learn by predicting one view in latent space from another and can emphasize semantic structure over low-level reconstruction [14–16]. This embedding-space principle has not yet been brought to single-cell transcriptomics.

Here we introduce CellOS, a generative multi-view single-cell foundation model that learns cellular representations from paired expression and perception views. CellOS keeps the cell-sentence interface but pairs the conventional expression view with a perception view that re-ranks genes by population-relative surprisal. The expression view answers what a cell expresses; the perception view asks which observations are least expected under the population and therefore most informative about identity, state or response. CellOS then aligns the two views with an LLM-JEPA objective, predicting agreement between the expression-view embedding and perception-view embedding while retaining causal language modelling so that generative structure is not discarded [15]. In this way, population-level salience becomes an explicit training signal rather than a heuristic hidden inside a single ranking.

CellOS is trained by a three-stage scaling recipe. In the first stage, a dense transformer learns the grammar of cellular expression from expression-view cell sentences. In the second, the converged dense model is expanded in a function-preserving manner into a mixture-of-experts transformer: the dense feed-forward weights seed one common expert, 32 cell-state-specific routed experts are introduced with near-zero initialization, and the converted model initially preserves the dense model’s behaviour while gaining capacity for continued pretraining. In the third stage, CellOS introduces the perception view and optimizes joint-embedding alignment together with the causal-language-modelling loss. A token-balanced data engine partitions distributed training by estimated token load, supporting pretraining of a 12-billion-parameter model on 390.5 million single-cell transcriptomes.

The contributions of CellOS are therefore threefold. First, it provides an information-theoretic complementary-view tokenization for cell sentences, separating expression magnitude from population-level information content so that rare but state-informative gene values are visible to the model. Second, it adapts joint-embedding predictive alignment to single-cell data; to our knowledge, this is the first application of JEPA-style representation learning to single-cell transcriptomics. Third, it couples this objective to a scalable dense-to-MoE training system, allowing capacity to grow without destabilizing a pretrained dense backbone. Together, these choices advance a representation-level thesis: a foundation model of the cell should encode not only what is expressed, but also which expression events are most informative in the population from which the cell arises.

## 2 Related Work

CellOS builds on three lines of work: single-cell foundation models, gene tokenization for cell-sentence learning and multi-view self-supervision. Together, these areas motivate CellOS’s central design: scale the cell-sentence paradigm, make population-level gene informativeness explicit and align complementary cellular views in latent space.

### 2.1 Foundation Models for Single-cell Transcriptomics

Single-cell foundation models emerged from earlier probabilistic models of cellular variation. scVI casts expression counts as observations from a latent-variable generative model and remains a standard baseline for denoising, batch correction and representation learning [17]. Transformer-based models then shifted the field from dataset-specific latent spaces toward reusable pretrained backbones. scBERT adapted masked language modelling to binned expression values for cell-type annotation [12]. Geneformer introduced a rank-ordered gene representation and showed that pretraining on approximately 30 million cells can support transfer to network-biology tasks with limited labelled data [4]. scGPT framed single-cell and multi-omic modelling as a generative pretraining problem [5], whereas scFoundation used an asymmetric encoder-decoder architecture to model raw transcriptomic measurements at large scale [6]. Related models such as UCE and scTab emphasize transferable cell embeddings and cross-tissue or cross-species generalization [18, 19].

Recent work has pushed both scale and ambition. TranscriptFormer trains a generative autoregressive model on 112 million cells across 12 species spanning 1.53 billion years of evolution; by coupling gene and transcript-count prediction with expression-aware attention, it recovers cell types, developmental trajectories and phylogenetic structure from unlabelled data [7]. State shifts the emphasis from observational atlases to perturbation response. It separates a State Embedding model pretrained on 167 million observational cells from a State Transition model trained on more than 100 million perturbed cells, using set-level self-attention to compare perturbed and control populations [20]. This direction is motivated by an important empirical warning: for perturbation-effect prediction, deep models have not always out-performed simple linear baselines [21]. Together with VCWorld and broader roadmaps for the AI virtual cell and perturbation cell atlas, these studies argue that the field is moving from cell annotation toward predictive, causal and queryable models of cellular state [8, 9, 22–24].

Despite their differences, most existing foundation models expose the model to one primary molecular view: expression counts, bins or rankings, trained with reconstruction, masked-modelling or autoregressive objectives. This is sufficient to learn many broad axes of variation, but it does not directly ask whether a gene’s value is informative relative to its own population distribution, nor whether two complementary views of the same cell should meet in a shared latent space. CellOS addresses this representational gap while preserving the scaling logic of prior foundation models: it adds a population-aware perception view, aligns it with the expression view and implements the training recipe on a dense-to-mixture-of-experts backbone.

### 2.2 Gene Tokenization and Cell-sentence Modelling

Tokenization determines which aspects of a transcriptome can be learned by a transformer. Current strategies fall into several families: rank-value encoding, as in Geneformer, sorts genes by normalized expression [4]; binning discretizes expression values before masked modelling, as in scBERT and scGPT [5, 12]; direct projection maps continuous counts into the model, as in scFoundation [6]; and gene-as-text or gene-embedding approaches represent genes through learned vectors, natural-language-derived embeddings or protein-sequence embeddings [7, 25, 26]. The most language-native representation is the cell sentence. Cell2Sentence renders each cell as a sequence of gene names ordered by expression, making transcriptomes accessible to large language models [10]; C2S-Scale extends this interface to much larger token corpora and model sizes [13].

These choices differ in implementation, but most remain organized around expression magnitude within a cell. Geneformer’s rank-value encoding is the strongest exception because it uses corpus-level normalization to reduce the dominance of ubiquitous genes and promote genes that better distinguish cellular state [4]. That design insight is important: the informativeness of a gene depends on its population context. However, a single normalized ranking still forces two questions into one order. It asks which genes are high in this cell after normalization, but not which gene values are statistically unexpected under each gene’s own distribution. For rare states, this distinction matters. A moderate expression value can be highly informative if it lies in the tail of the gene’s global distribution, whereas a large expression value may be biologically routine if most cells share it.

CellOS makes this population context explicit through a second token sequence. The expression view preserves the familiar magnitude-ranked cell sentence. The perception view re-ranks genes by population-relative surprisal estimated from each gene’s global expression distribution in the training corpus. This view is not intended to replace expression magnitude. Instead, it separates magnitude from salience, allowing the model to learn when the two agree and when they diverge. By treating population-level information content as a distinct intrinsic view of the same cell, CellOS turns a useful normalization principle into a trainable representation-learning objective.

### 2.3 Multi-view and Joint-embedding Representation Learning

Self-supervised learning increasingly distinguishes reconstruction of inputs from prediction in representation space. Contrastive approaches such as SimCLR, CLIP and contrastive predictive coding learn by bringing together embeddings of related views while separating unrelated examples [27–29]. Joint-embedding predictive architectures pursue a related but non-reconstructive principle: I-JEPA predicts embeddings of masked image regions from visible context, V-JEPA extends feature prediction to video, and data2vec provides a broader recipe for latent prediction across speech, vision and language [14, 30, 31]. The motivation is that a good representation should preserve the structure needed for prediction while ignoring nuisance details that raw reconstruction may overemphasize [16, 32]. LLM-JEPA adapts this idea to language models by adding a JEPA alignment term to next-token prediction, explicitly aligning different views in latent space rather than assuming that token prediction will align them automatically [15].

Single-cell biology has its own multi-view tradition, although it has usually taken a different form. scGen and CPA use latent spaces to model perturbation responses and covariates [33, 34]. GEARS combines gene-gene knowledge graphs with expression to predict transcriptional outcomes of unseen perturbations [35]. contrastiveVI separates salient variation from background variation in single-cell data [36]. State predicts perturbation effects in embedding space before decoding to expression, and VCWorld combines symbolic biological knowledge with data-driven modelling [20, 22]. These methods show that latent structure and auxiliary views can improve biological prediction, but they do not align two intrinsic tokenized views of the same transcriptome with a JEPA-style objective.

CellOS connects these threads. The expression and perception views are both derived from the same measured transcriptome, but they emphasize different statistical questions: abundance within the cell and information content under the population. Joint-embedding alignment forces these views to agree at the level of cell representation while causal language modelling preserves the sequential cell-sentence objective. This makes CellOS a biological analogue of latent-view prediction: the model is not only asked to reconstruct genes, but also to reconcile complementary meanings of the same cellular state.

## 3 Methodology

### 3.1 Overview of CellOS

CellOS implements an LLM-JEPA framework for single-cell transcriptomics, combining autoregressive cell-sentence modeling, sparse expert scaling and cross-view latent prediction. The input to CellOS is a collection of single-cell RNA-seq profiles, each represented over a shared gene vocabulary after gene-symbol harmonization and preprocessing. The pretraining corpus comprised 390,472,902 single-cell transcriptomes collected from 47,845 sequencing samples, of which approximately 346 million cells were associated with cell-level annotations. CellOS first converts each transcriptome into an expression-ranked gene sentence and trains a causal language model to capture the grammar of cellular expression. The dense language model is then expanded through sparse MoE scaling. A second, perception-ranked sentence is introduced to provide a complementary view of the same transcriptome for JEPA-based representation learning. The overall CellOS framework is illustrated in Fig. 1.

**Fig. 1.**
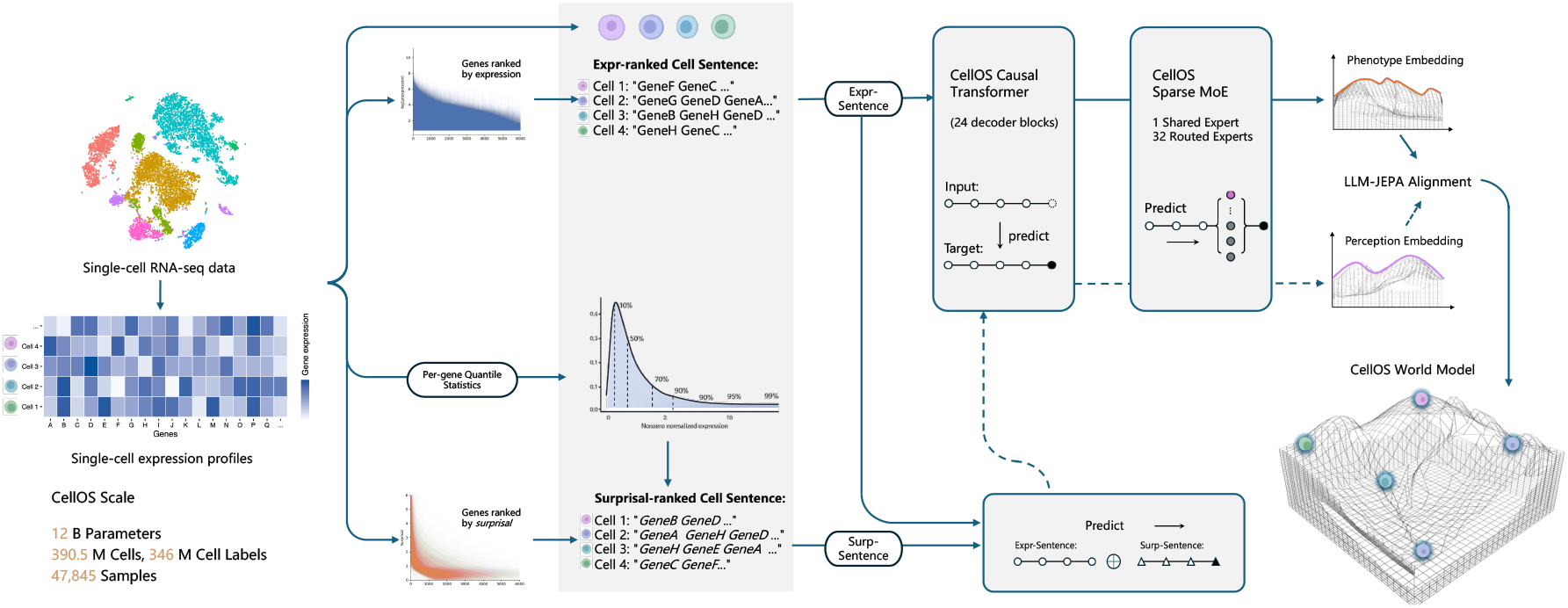
Overview of the CellOS framework. CellOS represents each single-cell transcriptome as complementary gene-token sentences. The expression-ranked sentence orders genes by normalized within-cell abundance, whereas the perception-ranked sentence orders genes by population-relative information content estimated from gene-specific expression statistics. The expression view is used to train a causal cell-sentence language model, which is subsequently expanded into a sparse mixture-of-experts Transformer. During the final stage, an LLM-JEPA objective aligns expression-view and perception-view representations in latent space, yielding unified cell embeddings for downstream analysis and prediction.

Both views are processed by a shared decoder-only causal Transformer. The expression view orders genes according to normalized expression abundance, whereas the perception view orders the same gene universe according to population-relative information content. The expression and perception views provide complementary descriptions of the same cellular state. Cross-view alignment is introduced through a joint-embedding predictive architecture in which an expression-view representation predicts the perception-view representation of the same cell in latent space. A dedicated representation token is appended to each cell sentence, and its final hidden state is used as the cell-level representation.

Training proceeds in three stages. First, a dense causal language model is pretrained on expression-view cell sentences. Second, dense feed-forward layers are converted into MoE layers by function-preserving initialization, followed by continued pretraining. Third, CellOS is trained with cross-view JEPA alignment while retaining autoregressive language modeling.

### 3.2 Data Processing and Gene Vocabulary Construction

CellOS operates on raw single-cell RNA-seq count matrices in which rows correspond to cells and columns correspond to genes. Let *x*_*cg*_ denote the raw count of gene *g* in cell *c*. Because count matrices originate from heterogeneous studies, protocols and genome annotations, gene identifiers are first harmonized to a common gene-symbol namespace. Symbols mapping to the same canonical gene are collapsed by summing their counts within each cell. Genes that cannot be resolved to the shared namespace or that fall outside the constructed vocabulary are excluded from sentence construction.

The gene vocabulary is formed from the harmonized gene universe observed across the pretraining corpus, together with reserved control tokens used for sequence delimitation and representation extraction. Each retained gene is assigned a unique token identifier. For a given cell, the count vector is mapped to this vocabulary and converted into a normalized expression profile. Library-size normalization is applied as

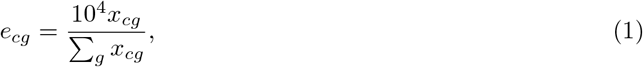

where *e*_*cg*_ denotes the normalized expression of gene *g* in cell *c*. This normalization places cells with different sequencing depths on a comparable scale before they are converted into rank-based sentences. It preserves relative within-cell abundance while reducing the direct influence of capture efficiency and library size on token order.

In addition to normalized expression values, CellOS estimates gene-specific empirical statistics over the pretraining population. For each gene, nonzero normalized expression values are summarized using empirical quantiles. These global statistics are computed from the harmonized training corpus and are used to define the perception view. The statistics are gene specific rather than cell type specific, allowing the perception view to express how unusual a normalized expression value is relative to the population distribution for the same gene.

### 3.3 Expression View: Expression-ranked Cell Sentences

The expression view represents each cell as a gene-token sentence ordered by decreasing normalized expression. For cell *c*, all retained genes with measured expression are sorted according to *e*_*cg*_. Formally, the ordered genes *g*_(1)_, *g*_(2)_, …, *g*_(*n*)_ satisfy

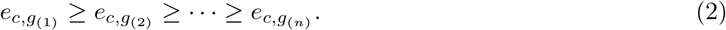

The expression-view sentence is then

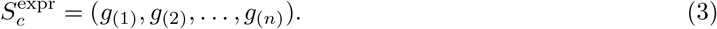

Ties are resolved deterministically to ensure reproducible tokenization. Reserved tokens are added according to the model input format, including a dedicated representation token from which the final cell embedding is extracted. This construction converts a sparse or high-dimensional expression vector into a discrete sequence while preserving the most salient within-cell abundance structure. Genes appearing early in 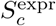 are those that dominate the measured transcriptome of the cell after normalization. The resulting sentence encodes the intrinsic transcriptional phenotype of the cell: highly abundant lineage markers, metabolic programs, structural components and other expressed genes are placed in positions where the autoregressive model is trained to assign high contextual probability.

The expression-ranked formulation follows the broad design principle used by Geneformer, Cell2Sentence and C2S-Scale, in which cells are transformed into ordered gene sequences suitable for language-model pretraining. In CellOS, this view provides the primary generative substrate for causal language modeling. It teaches the model regularities of cellular expression composition, including gene co-occurrence, shared sequence contexts and the organization of transcriptional programs within individual cells.

### 3.4 Perception View: Information-aware Cell Sentences

The perception view provides the complementary cellular signal used for JEPA alignment. We refer to this information-aware ordering as the perception view because it describes how a cell stands out relative to the statistical background of the population, rather than only what it abundantly expresses. A gene can be highly expressed because it is constitutively abundant across many cells, whereas a moderately expressed gene may carry high population-relative information content if its expression is unusually high relative to that gene’s own background. CellOS therefore constructs a second cell sentence in which genes are ordered by population-relative information content rather than by normalized abundance alone. The complementary-view formulation is illustrated in Fig. 1.

For each gene *g*, CellOS estimates an empirical quantile function from nonzero normalized expression values observed in the pretraining corpus. Given a cell *c*, the normalized expression value *e*_*cg*_ is mapped to a gene-specific empirical quantile *q*_*g*_(*e*_*cg*_). The perception information score is defined as the upper-tail quantile surprisal:

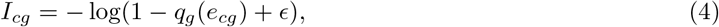

where *ϵ* is a small numerical constant. Larger values indicate that the normalized expression of gene *g* lies in an unusually high region of that gene’s empirical population distribution. The perception-view sentence 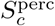 is obtained by ranking genes in cell *c* by decreasing *I*_*cg*_. Both views are constructed from the identical gene universe and differ only in their ranking criterion.

The score is quantile based rather than a continuous density estimate or a bin-frequency surprisal score. This choice is appropriate for single-cell expression, where values are sparse, overdispersed and strongly dependent on gene identity. Direct comparison of raw counts across genes would confound biological signal with sequencing depth and gene-specific abundance scale. Even after library-size normalization, genes differ markedly in dynamic range and prevalence. Gene-specific empirical quantiles place each normalized expression value in the context of the population behavior of the same gene, yielding a stable criterion across genes with different expression scales. The upper-tail form intentionally prioritizes genes whose expression is high relative to their own background.

The two views emphasize different aspects of cellular state. The expression view prioritizes transcriptional abundance, whereas the perception view prioritizes expression values that are elevated relative to the empirical population background of the same gene. Many single-cell signals are context dependent in this sense. Rare cellular states may be marked by a small set of genes that are not the most abundant transcripts in the cell but are elevated relative to their usual distribution. Transcription factors and signaling genes can have modest absolute abundance yet carry strong regulatory information when they appear in the upper tail of their gene-specific background. Stress-response genes and other inducible programs may become prominent because their expression departs from the population baseline. Context-specific programs, including transient activation, perturbation response and state transitions, are often defined by such relative deviations rather than by high abundance alone.

The two views provide complementary orderings of the same transcriptome. The expression view emphasizes the dominant molecular composition of the cell, whereas the perception view emphasizes upper-tail deviations from gene-specific population baselines. Processing both views through the same backbone provides the complementary inputs required for LLM-JEPA training, encouraging representations that capture both cellular identity and population-relative information content.

### 3.5 Model Architecture of CellOS

The architectural realization of CellOS is summarized in Fig. 2a–c, including complementary-view cell sentence construction, LLM-JEPA cross-view prediction and sparse MoE scaling. The CellOS architecture is organized around these three components. The expression and perception sentences provide complementary orderings of the same transcriptome. Both are processed by a shared decoder-only causal Transformer, denoted *f*_*θ*_, which maps a gene-token sequence to contextualized hidden states. A dedicated representation token is included in each sequence, and its final hidden state defines the cell-level representation used for cross-view prediction.

**Fig. 2.**
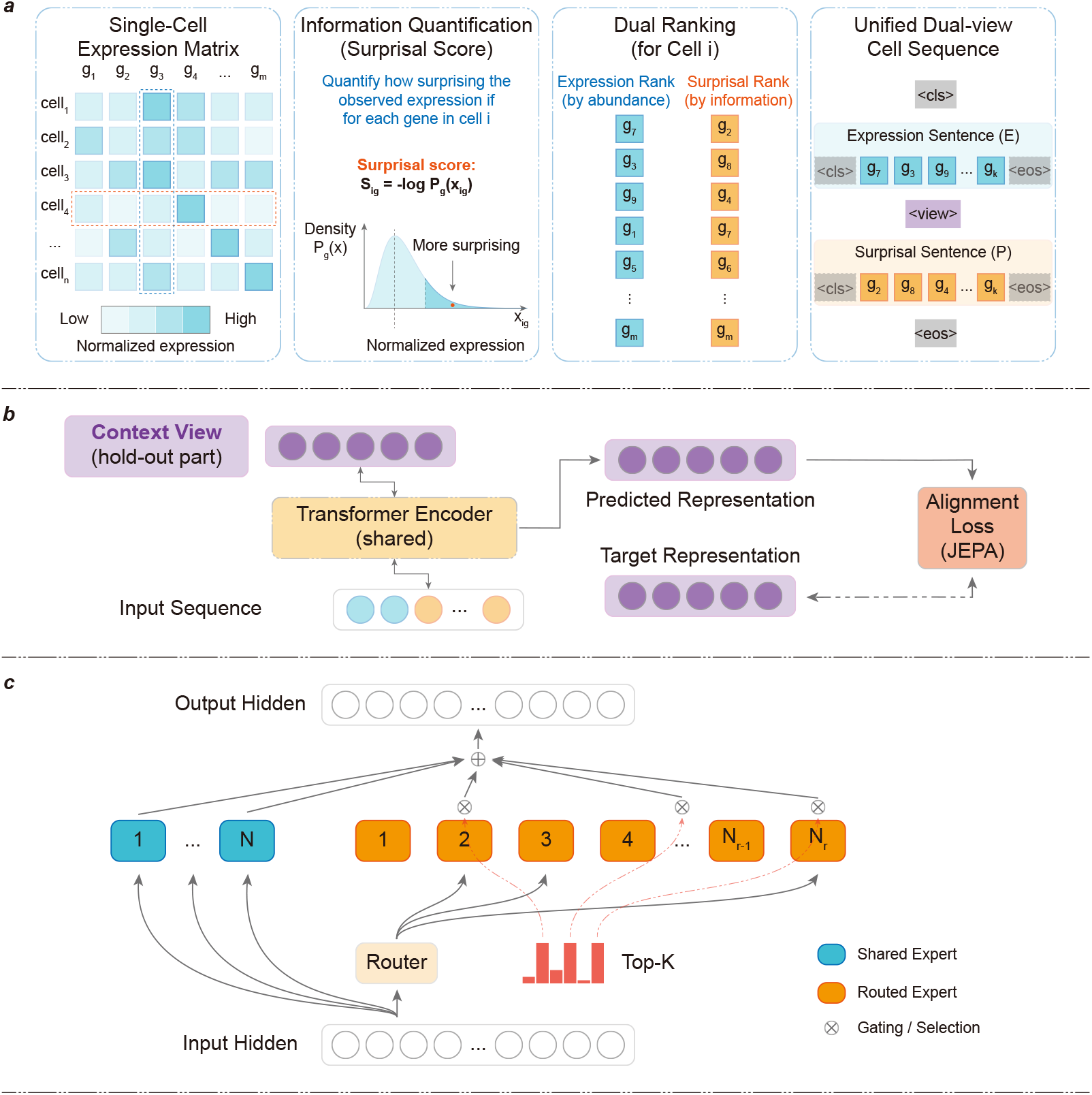
Architecture and training components of CellOS. **a**, Construction of expression-ranked and perception-ranked cell sentences from single-cell transcriptomes. The expression sentence is ordered by normalized abundance, whereas the perception sentence is ordered by population-relative information content. **b**, LLM-JEPA cross-view prediction aligns expression-view and perception-view representations in latent space. **c**, Dense-to-MoE scaling introduces shared and routed experts to expand model capacity while preserving a common computational pathway.

Autoregressive language modeling is applied to expression-ranked cell sentences to establish the cellular language model. Cross-view JEPA alignment then predicts the perception-view representation from the expression-view representation in latent space. Sparse MoE layers are introduced by dense-to-MoE conversion to expand model capacity during continued pretraining. Thus, the Transformer backbone serves as the shared sequence model, whereas the methodological focus of CellOS is the construction of biologically defined views, their latent alignment and scalable sparse capacity expansion.

#### 3.5.1 Causal Cell Sentence Modeling

Autoregressive language modeling is performed on expression-ranked cell sentences, as illustrated in Fig. 2a. Given 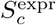, the model predicts each gene token from its preceding context, corresponding to *p*(*g*_*t*_ | *g*_<*t*_). Because the sentence order is determined by normalized expression, the prediction task exposes the model to recurrent patterns in within-cell transcriptional composition. Genes that appear in similar cellular contexts are therefore learned through their sequence neighborhoods and conditional dependencies.

This objective provides the generative foundation of CellOS. It trains the shared causal model to represent the grammar of cellular expression before the perception view is introduced. In this setting, “grammar” refers to the regular organization of genes within single-cell transcriptomes, including co-expression structure, lineage-associated expression programs and the ordering patterns induced by abundance-ranked cell sentences. The same language-modeling objective is retained during later stages so that sparse scaling and JEPA alignment are added without discarding the autoregressive cell-sentence modeling signal.

#### 3.5.2 JEPA Cross-view Prediction Module

The JEPA module is the principal representation-learning component of CellOS. The cross-view alignment mechanism is illustrated in Fig. 2b. For each cell, 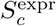 and 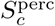 are constructed from the same transcriptome and the same gene vocabulary, but with different ranking criteria: normalized expression abundance for the expression view and gene-specific population-relative information content for the perception view. The two sequences therefore encode complementary descriptions of the same biological state.

This construction makes token-level reconstruction unsuitable. CellOS does not attempt to reconstruct one ranking from the other, because the two orderings are deliberately different. Such an objective would emphasize rank conversion rather than the cellular state shared across views. Instead, CellOS uses latent prediction to identify the aspects of the transcriptome that remain stable across abundance-driven and information-aware descriptions.

Each view is passed independently through the shared causal Transformer. The final hidden state of the representation token from the expression-view sequence defines 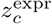, and the corresponding hidden state from the perception-view sequence defines 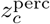. A predictor network *q*_*ϕ*_ maps the expression-view representation into the latent space of the perception view:

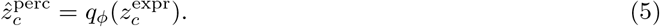

The alignment loss compares the predicted perception representation with the target perception representation using cosine distance:

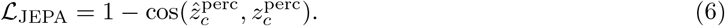

Unlike reconstruction-based objectives that recover genes, ranks or expression values in input space, CellOS predicts only latent representations. This avoids enforcing identical token sequences and directs learning toward cellular structure shared across the two views. The predictor is trained to recover the perception-view cell representation from the expression-view representation, encouraging the backbone to encode factors that are invariant across complementary rankings of the same transcriptome.

This formulation follows the joint-embedding predictive principle used in I-JEPA and the LLM-JEPA paradigm, where the target is an abstract representation of another view rather than the observed tokens themselves. In CellOS, the views are biologically defined rather than generated by generic masking or augmentation. Latent prediction therefore allows the model to integrate expression abundance with population-relative information content while preserving causal cell-sentence modeling as the foundation of the architecture.

The perception representation is treated as the target, and the predictor absorbs the transformation required to map expression-view information into the perception-view latent space. Because both views are processed by the same model, the JEPA objective regularizes the learned cell representation toward stability across complementary descriptions of the transcriptome. The resulting representation remains grounded in autoregressive expression modeling while incorporating population-relative information content not directly captured by expression magnitude.

#### 3.5.3 Mixture-of-Experts Scaling Module

CellOS expands the dense causal language model through a function-preserving dense-to-MoE conversion. The sparse scaling architecture is illustrated in Fig. 2c. After Stage 1 dense pretraining, selected feed-forward sublayers are replaced by MoE sublayers containing one shared expert and 32 routed experts. Routing is top-1: for each token, the router selects a single routed expert, while the shared expert remains active across tokens. This preserves a common computational pathway while adding conditional capacity.

The conversion is initialized to keep the sparse model close to the pretrained dense model. The shared expert is initialized from the corresponding dense feed-forward weights, routed experts are initialized near zero, and the router is initialized near uniform. These choices support stable transfer from the dense language model to the sparse model at the start of continued pretraining.

The MoE component provides scalable capacity expansion within the CellOS pipeline. It does not alter the cell-sentence formulation or the JEPA alignment objective. Continued pretraining allows the sparse model to adapt additional capacity to gene-token contexts, while MoE auxiliary losses regularize expert utilization.

### 3.6 Pre-training Objectives

CellOS combines three objectives during pretraining. The language modeling objective trains the model to predict gene tokens from expression-view contexts. The JEPA objective aligns expression-view and perception-view cell representations in latent space. The MoE auxiliary objectives regularize sparse routing during MoE pretraining. These objectives are introduced progressively across the three training stages so that the model first learns expression grammar, then expands capacity through sparse experts and finally integrates the perception view through cross-view prediction.

#### 3.6.1 Expression-token Prediction Loss

The expression-token prediction loss is the standard autoregressive cross-entropy loss over expression-view cell sentences. Given 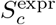, the model predicts each gene token from its preceding context under causal masking. Loss terms are accumulated over valid sequence positions and averaged over cells and tokens according to the training batch. This objective is used throughout all stages of pretraining. In Stage 1 it is the sole training signal, establishing the dense backbone. In later stages it preserves generative modeling of expression-ranked cellular sentences while MoE routing and JEPA alignment are added.

#### 3.6.2 Cross-view JEPA Alignment Loss

The cross-view JEPA alignment loss is applied to paired views of the same cell. The expression-view representation is passed through the predictor, and the predicted vector is aligned to the perception-view representation using the cosine-distance objective above. Alignment is performed in representation space rather than input space because the two views encode different orderings of the same gene vocabulary. Their token positions are not expected to match, and forcing token-level agreement would obscure the methodological role of the perception view. Latent alignment instead tests whether the expression view contains sufficient information to predict the cell-level semantics emphasized by the perception view. This encourages CellOS to form representations that integrate transcriptional abundance with population-relative information content.

#### 3.6.3 Multi-objective Training Strategy

The full training objective is

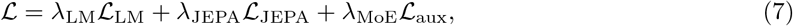

where the weights control the relative contribution of language modeling, cross-view alignment and MoE auxiliary regularization. In Stage 1, CellOS is trained as a dense causal language model using only the language modeling loss. In Stage 2, dense-to-MoE conversion is performed and continued pretraining uses the language modeling loss together with MoE auxiliary losses. In Stage 3, paired expression-view and perception-view sentences are introduced, and the model is optimized using language modeling, JEPA alignment and MoE auxiliary losses.

This staged optimization procedure integrates autoregressive language modeling, cross-view latent prediction and sparse expert scaling within a unified training framework.

## 4 Experiments

### 4.1 Experimental Settings

We evaluated CellOS on a unified single-cell benchmark covering both cell-state annotation and batch integration. We compared CellOS with six representative foundation-model baselines: UCE, State, scGPT, TranscriptFormer, STACK and C2S-2B. The annotation benchmark comprised six datasets spanning immune-cell annotation, developmental progression, fine-grained T-cell subclustering and perturbation-responsive cellular states, including PBMC immune aging, iPSC cell-type differentiation, iPSC developmental time-course, T-cell subclusters, human lung and IFN-*β*-stimulated PBMC datasets. Batch integration was evaluated on two datasets with explicit batch annotations: the human lung atlas and a perirhinal cortex dataset. Detailed dataset descriptions are provided in Supplementary Note 1.

Annotation performance was assessed using adjusted Rand index (ARI), normalized mutual information (NMI) and average silhouette width (ASW). For the aggregated benchmark shown in Fig. 3c, metrics were first normalized within each dataset and then averaged across datasets to obtain model-level annotation scores. Batch integration performance was evaluated using silhouette batch score (sil batch), graph connectivity (GC) and iLISI, which were averaged across the two batch datasets to obtain the aggregated batch-effect score. Detailed metric definitions and aggregation procedures are provided in Supplementary Note 2.

**Fig. 3.**
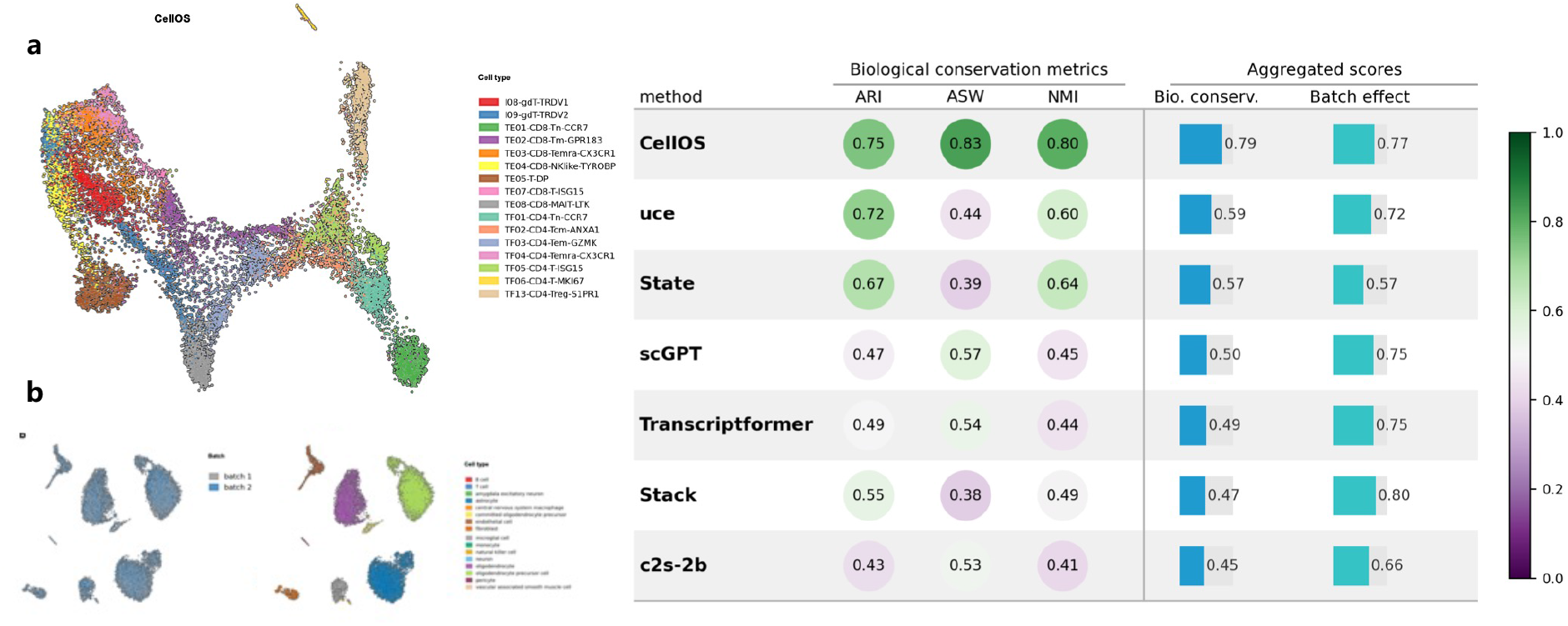
Evaluation of CellOS embeddings across annotation and integration tasks. **a**, UMAP visualizations of CellOS embeddings for a T-cell subcluster dataset colored by expert-annotated cell types, showing the organization of annotated cell states in the embedding space. **b**, UMAP visualizations of CellOS embeddings for the perirhinal cortex dataset colored by batch labels and cell types, presenting the same embeddings with respect to technical batch structure and biological identity. **c**, Average performance of CellOS and competing foundation models across annotation and batch-integration benchmarks. Annotation performance is summarized using dataset-normalized ARI, NMI and ASW scores, whereas integration performance is summarized using sil batch, graph connectivity (GC) and iLISI. Aggregated biological conservation and batch-effect scores are shown on the right.

To evaluate the contribution of model scaling and multi-view alignment, we further performed an ablation study on the T-cell subcluster benchmark using three CellOS checkpoints: 0.2B and 2B single-view pretrained models, and the final 12B MoE-scaled model trained with Stage-3 multi-view JEPA alignment.

### 4.2 CellOS Learns Generalizable Cell-state Representations

CellOS consistently outperformed existing foundation models across the aggregated annotation benchmark (Fig. 3c), achieving the highest normalized ARI (0.751), NMI (0.797) and ASW (0.828), and consequently the highest overall biological conservation score (0.792). This pattern was also evident on the lung atlas, where CellOS achieved strong raw annotation performance (ARI = 0.769, NMI = 0.858, ASW = 0.631), and remained competitive on PBMC and T-cell subcluster benchmarks. Together, these results indicate that CellOS preserves cell-state structure across diverse tissues, label granularities and perturbation contexts.

CellOS also maintained a favourable trade-off between biological conservation and batch mixing. In the batch benchmark (Fig. 3b,c), it achieved the highest graph connectivity (GC = 0.964) and a strong batch-effect score (0.771), second only to STACK (0.801). However, STACK showed substantially weaker biological conservation (0.474 versus 0.792 for CellOS). These results suggest that CellOS improves integration while retaining biologically meaningful neighborhood structure, rather than achieving batch mixing through over-integration.

### 4.3 Perturbation-response Prediction

We next asked whether CellOS embeddings provide a stronger substrate for perturbation modeling than existing single-cell foundation models.

Perturbation-response prediction was evaluated using the benchmark and transition-model framework introduced by STATE. To isolate the contribution of representation quality, only the foundation-model embeddings were varied across experiments, whereas the downstream transition model, training procedure and evaluation protocol were kept identical for all methods. The benchmark comprised five held-out cellular contexts (H1, HepG2, Jurkat, K562 and RPE1), and detailed dataset descriptions and evaluation procedures are provided in Supplementary Note 3.

As summarized in Fig. 4, CellOS consistently occupied the preferred region with both high perturbation fidelity and high biological consistency. Detailed quantitative results are reported in Table 1. CellOS achieved the strongest overall performance, ranking first on DE Spearman, Pearson*e*dist, clustering agreement and DE direction matching (Table 1). Notably, CellOS obtained the highest DE Spearman score (0.590), indicating the most accurate recovery of perturbation-induced differential-expression programs. CellOS also substantially outperformed all competing embeddings in Pearson*e*dist (0.619 versus 0.373 for the next-best model), suggesting that predicted perturbation responses more faithfully preserved global transcriptomic state transitions. In addition, CellOS achieved the highest clustering agreement (0.633), indicating improved recovery of perturbation-induced cellular organization at the population level. CellOS also matched the best-performing model on DE direction matching, further demonstrating that its learned representations effectively capture perturbation-induced transcriptional changes. Overall, CellOS consistently provided the most informative representations for downstream perturbation modeling across diverse evaluation criteria.

**Table 1.** Perturbation-response prediction across foundation-model embeddings. Reported values are means across H1, HepG2, Jurkat, K562 and RPE1 held-out test contexts. Higher values indicate better performance. Best values are shown in bold and second-best values are underlined.

| Model | DE Spearman<sub>sig</sub> | $\text{Pearson}_{\text{edist}}$ | Clustering agreement | DE direction match |
| --- | --- | --- | --- | --- |
| CellOS | **0.590** | **0.619** | **0.633** | **0.623** |
| TranscriptFormer | <u>0.582</u> | <u>0.373</u> | 0.611 | 0.520 |
| UCE | -0.023 | -0.113 | <u>0.625</u> | **0.623** |
| C2S | 0.463 | 0.124 | 0.618 | 0.608 |
| scGPT | 0.034 | 0.067 | 0.612 | 0.592 |
| STACK | 0.391 | 0.217 | 0.606 | 0.582 |
| State | -0.071 | -0.108 | 0.621 | 0.563 |

**Fig. 4.**
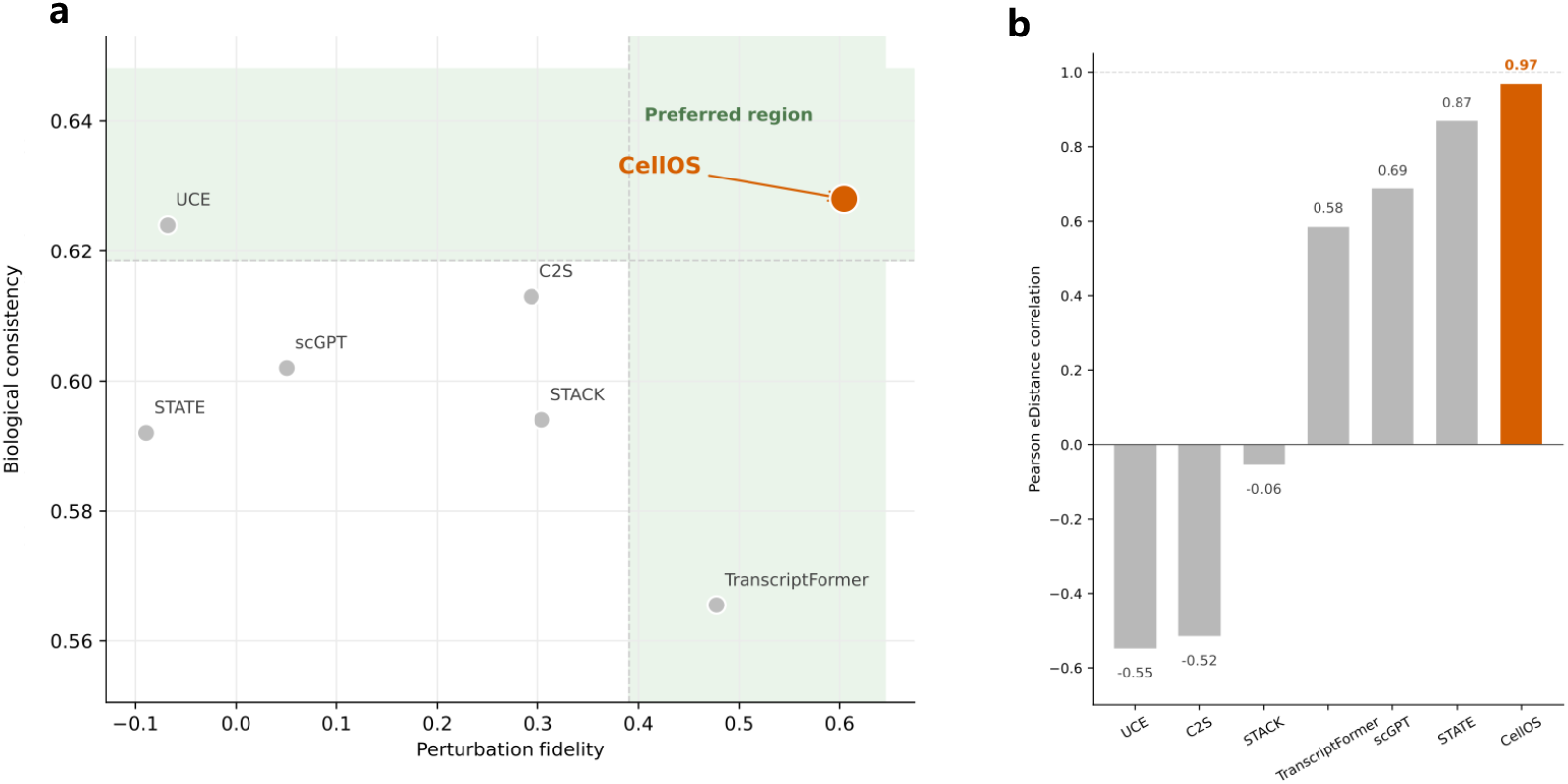
CellOS improves perturbation-response prediction across complementary evaluation metrics. **a**, Joint comparison of perturbation fidelity and biological consistency across foundation-model embeddings. Perturbation fidelity is defined as the mean of DE Spearman and Pearson_*e*dist_, whereas biological consistency is defined as the mean of clustering agreement and DE direction matching. The shaded upper-right region indicates the preferred operating regime with high performance on both criteria. **b**, Pearson_*e*dist_ comparison on the Jurkat held-out context. CellOS achieves the highest Pearson_*e*dist_ in this context, indicating improved preservation of global transcriptomic state transitions following perturbation.

Together, Fig. 4 and Table 1 show that the advantages of CellOS extend beyond individual metrics. These results suggest that the representations learned by CellOS capture biologically meaningful state variables that extend beyond static cell identity. Whereas annotation benchmarks primarily assess the separation of existing cellular states, perturbation prediction requires representations that preserve how cellular states change under genetic intervention. The strong performance of CellOS in this setting therefore indicates that the complementary expression and perception views encode information relevant not only to cellular identity but also to cellular response. This observation is consistent with the central design principle of CellOS: integrating transcriptional abundance with population-relative information content yields representations that remain informative under cellular state transitions, providing a stronger foundation for predictive models of cellular behavior.

### 4.4 Scaling and Multi-view Alignment Enhance Fine-grained Cell-state Resolution

On the T-cell subcluster benchmark, annotation performance improved systematically with both model scale and training stage (Fig. 5a). The 0.2B and 2B single-view pretraining variants achieved ARI/NMI/ASW of 0.395/0.579/0.524 and 0.497/0.660/0.523, respectively, whereas the 12B MoE-scaled Stage-3 checkpoint reached 0.539/0.688/0.531 (biological conservation score = 0.586). ARI and NMI increased monotonically with model scale (+36% and +19% relative to 0.2B), following an approximately log-linear trend with parameter count, whereas ASW remained near 0.52–0.53 across checkpoints. Scaling from 0.2B to 2B accounted for most of the gain (ΔARI = +0.102; ΔNMI = +0.081), with further improvements from the 12B MoE-scaled Stage-3 model (ΔARI = +0.042; ΔNMI = +0.028). Together, these results indicate that CellOS exhibits favorable scaling behavior with model size, while the additional gains observed in the Stage-3 checkpoint suggest that multi-view JEPA alignment provides benefits beyond parameter scaling alone. This scaling trend accompanies the expansion of CellOS to 12B parameters, positioning it among the largest single-cell foundation models compared in Fig. 5b.

**Fig. 5.**
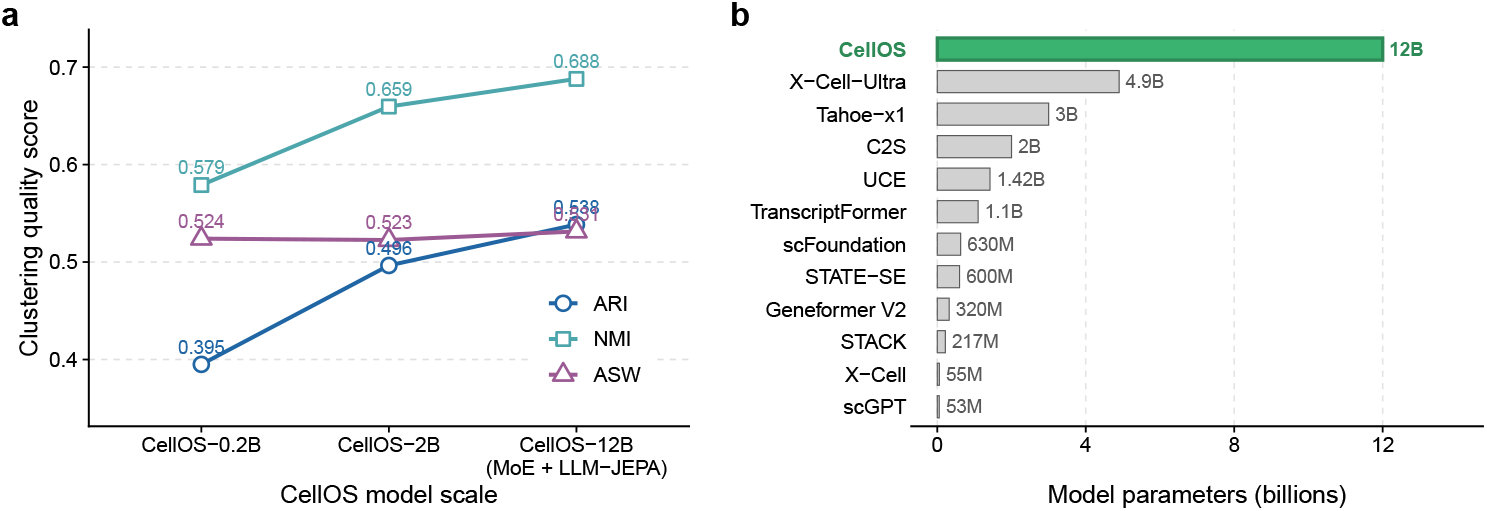
Scaling behavior of CellOS and model-size comparison with existing foundation models. **a**, ARI, NMI and ASW scores on the T-cell subcluster benchmark across CellOS checkpoints of increasing scale. **b**, Parameter-scale comparison between CellOS and representative single-cell foundation models. CellOS reaches 12B parameters through sparse MoE scaling, making it one of the largest single-cell foundation models reported to date.

## 5 Discussion

We developed CellOS, a multi-view foundation model for single-cell transcriptomes. CellOS reframes the cell-sentence paradigm by representing each transcriptome through two complementary views: an expression view that captures within-cell abundance and a perception view that captures population-relative surprisal. This design addresses a central limitation of expression-ranked cell sentences: highly expressed genes are not always the most informative genes for defining cellular state. By separating abundance from population-aware informativeness, CellOS provides a representation in which both dominant transcriptional programs and rare, context-dependent signals can contribute to the learned cell embedding.

Across annotation, batch-integration and perturbation-response benchmarks, CellOS produced transferable representations of cellular state. Its strong annotation performance indicates that CellOS preserves discrete biological identities across tissues and label granularities, whereas its integration results suggest that this preservation is not achieved at the expense of technical robustness. More importantly, CellOS embeddings also improved perturbation-response prediction when used within a shared State Transition framework. Because perturbation prediction requires representations that remain informative under state changes, these results suggest that CellOS captures structure beyond static cell-type identity and encodes latent variables relevant to cellular transitions.

A key implication of CellOS is the shift from reconstruction-centric learning toward representation-centric learning in single-cell foundation models. Most existing models learn by reconstructing tokens, expression bins or masked values. CellOS instead asks whether two biologically defined views of the same transcriptome can be aligned in latent space. The perception view is not an alternative measurement, but a second interpretation of the same cellular state. LLM-JEPA alignment encourages the model to recover information that remains stable across these complementary interpretations, rather than simply reproducing one input format. In this sense, CellOS follows the broader movement in self-supervised learning from low-level reconstruction toward predictive latent representation learning.

This perspective also connects CellOS to the idea of a cellular world model. In artificial intelligence, world models aim to learn latent states that explain multiple observations of the same underlying system. In CellOS, expression-ranked and perception-ranked sentences act as two observations of the same cell. Their agreement in latent space provides a route to learning a shared cellular state representation that is not tied to a single ranking rule. Although CellOS does not yet model full cellular dynamics, its transfer to perturbation-response prediction suggests that multi-view transcriptomic pretraining can provide a foundation for future models that reason about cellular state transitions, interventions and responses.

## Limitations and Future Work

Several limitations remain. First, the current perception view is estimated from gene-specific population statistics over the full pretraining corpus and does not explicitly model tissue-, lineage- or condition-specific backgrounds. Future work could introduce context-aware perception views that define informativeness relative to specific biological environments. Second, CellOS currently uses transcriptomic measurements alone. Incorporating chromatin accessibility, protein abundance, spatial information and perturbation readouts could provide additional views of cellular state and further improve representation learning. Finally, latent agreement between complementary views does not by itself establish a causal cellular model. CellOS learns representations that support downstream perturbation prediction, but it is not pretrained directly on perturbation trajectories. Extending CellOS with temporal and perturbational data will be important for moving from cellular representation learning toward predictive virtual-cell systems.

## Supplementary Information

### Supplementary Note 1. Benchmark Datasets

#### PBMC immune aging dataset

For the cell-type annotation benchmark, we analyzed a processed PBMC dataset derived from the Immune Aging Project (Wells et al., 2025; https://doi.org/10.1038/s41590-025-02241-4). After quality control and subsampling, the dataset comprised 29,669 cells and 25,071 genes from 14 tissues, including spleen, lymph nodes, blood, lung and skin. Cells were annotated into 13 cell types using the Scelltype label, including CD8 T, Memory B, Memory CD4, Monocyte and NK dim cells. Per-type counts ranged from 1,649 to 2,400 cells. This benchmark evaluated whether embeddings preserved expert-defined immune cell identities.

#### iPSC cell-type differentiation dataset

A processed iPSC dataset from GSE279710 was included to assess embedding performance on lineage annotation. The final object contained 9,029 cells and 25,525 genes, annotated into 25 differentiation states using the Annotation label, including Mesendoderm, Definitive endoderm, Posterior foregut and Pancreas. Cell-type counts ranged from 20 to 1,539 cells per class. Although the original study included multiple sequencing runs, four technical batches (run: 0Xav, lib1, lib2 and lib3) were retained for reference only; the annotation task was based on cell-type labels rather than batch labels.

#### iPSC differentiation time-course dataset

To evaluate whether embeddings captured temporal progression during iPSC differentiation, we included a processed dataset from the study described in PMC7010688. After processing, the dataset contained 36,044 cells, 11,231 genes and 125 donors. Cells were labeled by differentiation day, including day0, day1, day2 and day3, with 8,455 to 9,661 cells per time point.

#### T-cell subcluster dataset

Fine-grained T-cell subtype annotation was assessed using a processed dataset from GSE254249. The object contained 14,260 cells and 1,578 genes from 24 patients, sampled from a single tissue context across four time points using the SampleTimePoint label. Expert annotations defined 16 T-cell subclusters using the subCluster label, including I08-gdT-TRDV1 and TE01-CD8-Tn-CCR7. Most subclusters contained approximately 1,000 cells, with a range of 122 to 1,000 cells.

#### Human lung dataset

A processed human lung dataset based on Adams et al. (2020) was used for both annotation and batch-effect evaluation. The object comprised 46,091 cells, 29,604 genes and 107 sequencing libraries using the Library Identity label, corresponding to 75 libraries after merging by leading library identifier. Expert annotations defined 16 lung cell types using the Scelltype label, including Macrophage, cMonocyte, NK, Ciliated and B cells, with 567 to 7,644 cells per type.

For the annotation task, UMAP visualization and quantitative metrics were computed using Scelltype labels. For the batch-effect task, library identity was used to assess cross-library mixing while preserving cell-type structure.

#### IFN-*β*-stimulated PBMC dataset

The annotation benchmark also included a processed PBMC dataset from GSE96583 (Kang et al., 2018). After processing, the dataset contained 13,999 cells and 11,784 genes, split into IFN-*β*-stimulated cells (STIM, 7,451 cells) and control cells (CTRL, 6,548 cells). Unsupervised clustering yielded 13 groups using the seurat clusters label, with cluster sizes ranging from 56 to 4,321 cells. In this setting, evaluation was performed using cluster identity rather than stimulation condition.

#### Perirhinal cortex dataset

Batch-effect robustness was further examined using a processed perirhinal cortex dataset from Siletti et al. (Science 382, eadd7046, 2023; https://doi.org/10.1126/science.add7046). The object contained 17,535 cells and 58,232 genes from Brodmann area 35, split across two samples: 10X222 1 with 8,465 cells and 10X222 2 with 9,070 cells. Original annotations defined 16 cell types using the cell type label, including oligodendrocyte precursor cell, astrocyte and oligodendrocyte. For the batch-effect benchmark, UMAP visualization and metrics were computed using sample identity as the batch label, while cell-type annotations were used separately to assess preservation of biological structure alongside batch mixing.

### Supplementary Note 2. Evaluation Metrics and Aggregated Benchmark Construction

Annotation performance was evaluated using adjusted Rand index (ARI), normalized mutual information (NMI) and average silhouette width (ASW). Given reference labels **y** and predicted cluster assignments **ŷ**, ARI was computed as

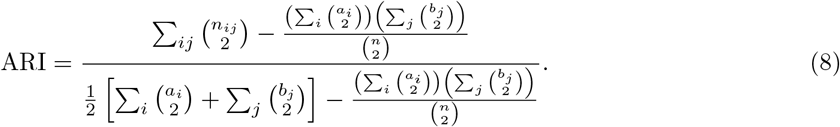

NMI was computed as

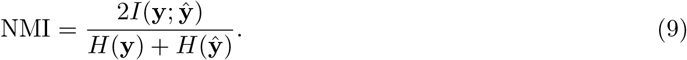

ASW was computed from the silhouette coefficient

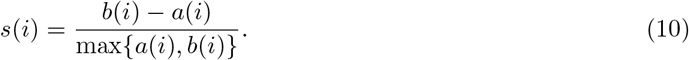

The dataset-level ASW was defined as

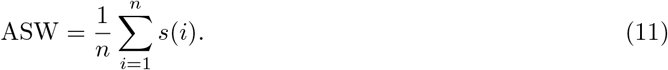

For aggregated annotation benchmarking, ARI, NMI and ASW were first normalized within each dataset using min-max normalization:

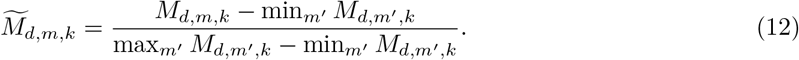

Normalized scores were averaged across the six annotation datasets. The aggregated biological conservation score was defined as

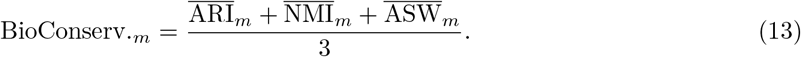

Batch integration was evaluated using silhouette batch score (sil batch), graph connectivity (GC) and integration local inverse Simpson’s index (iLISI). GC was defined as

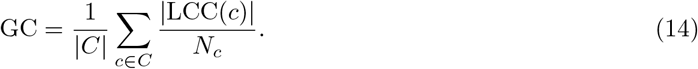

The iLISI metric was defined as

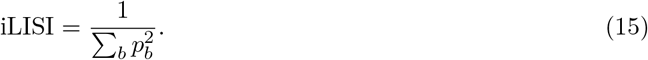

Metrics were averaged across the two batch-evaluation datasets. The aggregated batch-effect score was defined as

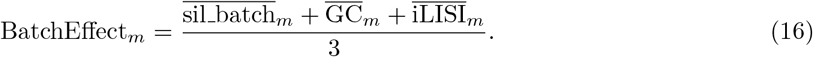

### Supplementary Note 3. Perturbation-response Prediction Benchmark

To evaluate whether CellOS embeddings capture perturbation-relevant cellular state information, we adopted the perturbation-response prediction framework introduced by STATE. In this benchmark, foundation-model embeddings are used as inputs to a shared perturbation transition model, allowing the contribution of representation quality to be evaluated independently from downstream model architecture. The benchmark was constructed from publicly available Perturb-seq datasets provided through the Virtual Cell Challenge resource. We used five commonly studied cellular contexts: H1 human embryonic stem cells (H1), HepG2, Jurkat, K562 and RPE1. These datasets comprise large-scale single-cell genetic perturbation experiments in which transcriptomic responses were measured following targeted gene perturbations. Together, they span pluripotent, epithelial and hematopoietic cellular states and therefore provide a diverse evaluation setting for perturbation generalization. (Virtual Cell Challenge)

For all compared methods, the downstream perturbation transition model, training procedure and evaluation protocol were kept identical. Only the foundation-model embeddings were replaced. Specifically, embeddings generated by CellOS, UCE, State, scGPT, TranscriptFormer, STACK and C2S were separately provided as inputs to the same transition model. This design isolates the effect of representation quality from differences in downstream model architecture.

Performance was evaluated using the metrics reported by the original STATE framework, including differential-expression recovery, transcriptomic-distance preservation and perturbation-induced clustering consistency. Results reported in Table 1 correspond to averages across the five held-out cellular contexts.

## Notes

### Competing Interest Statement

The authors are affiliated with Vitaura (Beijing) Technology Co., Ltd. A patent application related to the CellOS technology described in this manuscript is pending. The authors declare no other competing interests.

### Summary of Updates

This revised version includes updates to the abstract and manuscript text for improved clarity and readability. We also added a new figure to further illustrate the benchmarking results and updated the presentation of experimental results. These revisions improve the presentation of the work without affecting the main conclusions of the study.

## References

[1] Svensson, V., Vento-Tormo, R., Teichmann, S.A.: Exponential scaling of single-cell RNA-seq in the past decade. Nature Protocols 13, 599–604 (2018) 10.1038/nprot.2017.149

[2] Rood, J.E., et al.: The Human Cell Atlas: from a cell census to a unified foundation model. Nature 637, 1065–1071 (2025) 10.1038/s41586-024-08338-4

[3] CZI Cell Science Program: CZ CELLxGENE discover: a single-cell data platform for scalable exploration, analysis and modeling of aggregated data. Nucleic Acids Research 53, 886–900 (2025) 10.1093/nar/gkae1142

[4] Theodoris, C.V., et al.: Transfer learning enables predictions in network biology. Nature 618, 616–624 (2023) 10.1038/s41586-023-06139-9

[5] Cui, H., et al.: scGPT: toward building a foundation model for single-cell multi-omics using generative ai. Nature Methods 21, 1470–1480 (2024) 10.1038/s41592-024-02201-0

[6] Hao, M., et al.: Large-scale foundation model on single-cell transcriptomics. Nature Methods 21, 1481–1491 (2024) 10.1038/s41592-024-02305-7

[7] Pearce, J.D., et al.: TranscriptFormer: a generative cell atlas across 1.5 billion years of evolution. Science (2026) 10.1101/2025.04.25.650731

[8] Bunne, C., et al.: How to build the virtual cell with artificial intelligence: priorities and opportunities. Cell 187, 7045–7063 (2024) 10.1016/j.cell.2024.11.015

[9] Rood, J.E., Hupalowska, A., Regev, A.: Toward a foundation model of causal cell and tissue biology with a perturbation cell and tissue atlas. Cell 187, 4520–4545 (2024) 10.1016/j.cell.2024.07.035

[10] Levine, D., et al.: Cell2Sentence: teaching large language models the language of biology. In: Proceedings of the 41st International Conference on Machine Learning (ICML), PMLR, vol. 235, pp. 27299–27325 (2024). 10.1101/2023.09.11.557287

[11] Shen, H., et al.: Generative pretraining from large-scale transcriptomes for single-cell deciphering. iScience 26, 106536 (2023) 10.1016/j.isci.2023.106536

[12] Yang, F., et al.: scBERT as a large-scale pretrained deep language model for cell type annotation of single-cell RNA-seq data. Nature Machine Intelligence 4, 852–866 (2022) 10.1038/s42256-022-00534-z

[13] Rizvi, S.A., et al.: Scaling large language models for next-generation single-cell analysis (Cell2Sentence-Scale). bioRxiv 2025.04.14.648850 (2025). 10.1101/2025.04.14.648850

[14] Assran, M., et al.: Self-supervised learning from images with a joint-embedding predictive architecture (I-JEPA). In: Proceedings of the IEEE/CVF Conference on Computer Vision and Pattern Recognition (CVPR), pp. 15619–15629 (2023). arXiv:2301.08243

[15] Huang, H., LeCun, Y., Balestriero, R.: LLM-JEPA: large language models meet joint embedding predictive architectures. arXiv:2509.14252 (2025)

[16] Balestriero, R., LeCun, Y.: Learning by reconstruction produces uninformative features for perception. arXiv:2402.11337 (2024)

[17] Lopez, R., Regier, J., Cole, M.B., Jordan, M.I., Yosef, N.: Deep generative modeling for single-cell transcriptomics. Nature Methods 15, 1053–1058 (2018) 10.1038/s41592-018-0229-2

[18] Rosen, Y., et al.: Universal Cell Embeddings: a foundation model for cell biology. bioRxiv 2023.11.28.568918 (2023). 10.1101/2023.11.28.568918

[19] Fischer, F., et al.: scTab: scaling cross-tissue single-cell annotation models. Nature Communications 15, 6611 (2024) 10.1038/s41467-024-51059-5

[20] Adduri, A., et al.: Predicting cellular responses to perturbation across diverse contexts with State. bioRxiv 2025.06.26.661135 (2025). 10.1101/2025.06.26.661135

[21] Ahlmann-Eltze, C., Huber, W., Anders, S.: Deep-learning-based gene perturbation effect prediction does not yet outperform simple linear baselines. Nature Methods 22, 1657–1661 (2025) 10.1038/s41592-025-02772-6

[22] Wei, Z., et al.: VCWorld: a biological world model for virtual cell simulation. arXiv:2512.00306; ICLR 2026 (2025)

[23] Zhang, J., et al.: Tahoe-100M: a giga-scale single-cell perturbation atlas for context-dependent gene function and cellular modeling. bioRxiv 2025.02.20.639398 (2025). 10.1101/2025.02.20.639398

[24] Replogle, J.M., et al.: Mapping information-rich genotype-phenotype landscapes with genome-scale Perturb-seq. Cell 185, 2559–2575 (2022) 10.1016/j.cell.2022.05.013

[25] Guo, F., et al.: Foundation models in bioinformatics. National Science Review 12, 028 (2025) 10.1093/nsr/nwaf028

[26] Chen, Y.T., Zou, J.: GenePT: a simple but effective foundation model for genes and cells built from ChatGPT. bioRxiv 2023.10.16.562533 (2023). 10.1101/2023.10.16.562533

[27] Chen, T., Kornblith, S., Norouzi, M., Hinton, G.: A simple framework for contrastive learning of visual representations (SimCLR). In: Proceedings of the 37th International Conference on Machine Learning (ICML), PMLR, vol. 119, pp. 1597–1607 (2020)

[28] Radford, A., et al.: Learning transferable visual models from natural language supervision (CLIP). In: Proceedings of the 38th International Conference on Machine Learning (ICML), PMLR, vol. 139,pp. 8748–8763 (2021)

[29] Oord, A., Li, Y., Vinyals, O.: Representation learning with contrastive predictive coding. arXiv:1807.03748 (2018)

[30] Bardes, A., et al.: Revisiting feature prediction for learning visual representations from video (V-JEPA). arXiv:2404.08471 (2024)

[31] Baevski, A., Hsu, W.-N., Xu, Q., Babu, A., Gu, J., Auli, M.: data2vec: a general framework for self-supervised learning in speech, vision and language. In: Proceedings of the 39th International Conference on Machine Learning (ICML), PMLR, vol. 162, pp. 1298–1312 (2022). arXiv:2202.03555

[32] LeCun, Y.: A path towards autonomous machine intelligence. OpenReview (2022)

[33] Lotfollahi, M., Wolf, F.A., Theis, F.J.: scGen predicts single-cell perturbation responses. Nature Methods 16, 715–721 (2019) 10.1038/s41592-019-0494-8

[34] Lotfollahi, M., et al.: Predicting cellular responses to complex perturbations in high-throughput screens. Molecular Systems Biology 19, 11517 (2023) 10.15252/msb.202211517

[35] Roohani, Y., Huang, K., Leskovec, J.: Predicting transcriptional outcomes of novel multigene perturbations with GEARS. Nature Biotechnology 42, 927–935 (2024) 10.1038/s41587-023-01905-6

[36] Weinberger, E., Lin, C., Lee, S.-I.: Isolating salient variations of interest in single-cell data with contrastiveVI. Nature Methods 20, 1336–1345 (2023) 10.1038/s41592-023-01955-3

